# Principled measures and estimates of trait polygenicity

**DOI:** 10.1101/2025.07.10.664154

**Authors:** Luke J. O’Connor, Guy Sella

## Abstract

The ‘polygenicity’ of traits is often invoked and sometimes quantified in quantitative, statistical, and human genetics. What do we mean by the polygenicity of a trait? We propose a principled definition that encompasses a range of polygenicity measures. We show that these measures satisfy certain mathematical properties, we argue that these properties are sensible if not necessary, and we show that, conversely, measures that satisfy these properties also satisfy our definition. We consider four specific measures in greater detail, describe how they differ and show that three of them can be estimated from GWAS summary statistics using an existing method, Fourier Mixture Regression. We estimate these measures for 36 traits in humans. We find a dearth of traits with polygenicity values that fall within the large gap between Mendelian and highly polygenic traits. We discuss the evolutionary and cellular processes underlying trait polygenicity.

## Introduction

The ‘complexity’ or ‘polygenicity’ of traits varies immensely. In one extreme, there are Mendelian or simple traits with most heritable variance arising from variants in a single gene. A few are common in specific human populations or geographic regions (e.g., sickle cell anemia), and many are rare^1^. We focus on examples from humans, which we typically know more about, but examples abound in many plants and animals.

In the other extreme, there are complex or highly polygenic traits. Examples include morphological traits (e.g., height^2^), life history traits, physiological traits, and biomarkers, as well as the risk of most common genetic diseases (e.g., schizophrenia^3^). Here, too, examples from other taxa abound.

In quantitative genetics, complex traits were assumed to be highly polygenic for nearly a century^4^, but the degree of polygenicity has become evident only recently, owing to genome-wide association studies (GWAS) in huge samples^5^ performed in humans. These studies revealed that many traits are much more polygenic than previously appreciated (see, e.g., ref. ^6^), with heritable variance thinly spread among myriad variants throughout the genome^2,3,7–12^. Estimates of the number of common variants that contribute to heritable variation in Europeans range from several hundred to over a hundred thousand, across traits and estimation methods^13–17^.

Measuring polygenicity in terms of the number of variants with nonzero contributions to heritable variance, however, raises both conceptual and statistical problems. The statistical challenge is to account for variants with arbitrarily small (nonzero) contributions. Current estimation approaches model the distribution of contributions to variance as a mixture of a point mass at 0 with a single Normal distribution^13,18,19^ or 2-3 Normal distributions ^14,20^. Violations of these modeling assumptions plausibly lead to substantial downward biases of the polygenicity estimate^14,21^.

The conceptual problem with this measure of polygenicity is arguably more severe. Suppose that variation in one trait arises from 10,000 causal variants with equal contributions to variance, whereas variation in another trait arises from one large-effect variant that contributes 99% of the variance alongside 9,999 variants that contribute the remaining 1%. We would consider the first trait to be highly polygenic and the second to be nearly Mendelian, yet the commonly assumed measure would assign a polygenicity of 10,000 to both.

The ‘effective’ polygenicity measure proposed by O’Connor and colleagues circumvents these problems by downweighting the contributions to polygenicity of variants with small contributions to variance^21^. In our example, this measure would assign a polygenicity of 10,000 to the first trait and ∼1.01 to the second. The weighting eases estimation because it makes little difference to the estimate whether an effect is zero vs. nearly-zero. This polygenicity measure, however, corresponds to a particular choice about how to take an average across variants (see below), raising the question of whether there are alternatives, possibly more suitable ones, and more generally, how polygenicity should be measured.

Here we propose a general definition of polygenicity measures, which matches our intuitions and has sensible mathematical properties. We consider several measures of interest that satisfy this definition, analyze the relationship among them, and estimate three of them for 36 human traits.

## Methods

### Three measures of polygenicity

We first consider polygenicity measures in a simple setting. We assume that additive heritable variation in a trait in a given population arises from *n* biallelic variants that are at linkage equilibrium (LE) with one another (we consider linkage disequilibrium later). We denote the proportional contribution of variant *i* to additive variance by *p*_*i*_ and the distribution of variance over all variants by *D* = (*p*_1_, …, *p*_*n*_), where *p*_*i*_ > 0 for *i* = 1, …, *n*, and 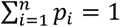. In this setting, a measure of polygenicity is a function of the distribution of additive genetic variance across variants, i.e., *P* = *P* (*D*).

There is one special case in which we know what value this function should take. When additive variance is equally distributed among *n* variants, i.e., when *p*_1_ = *p*_2_ = ⋯ = *p*_*n*_ = 1⁄*n*, we would like the polygenicity to be equal to *n*.

This special case suggests how we may attribute a polygenicity to any given variant. We define the polygenicity of non-null variant *i* as π_*i*_ ≡ 1⁄*p*_*i*_. This is a *variant polygenicity* in the sense that a trait whose variants all contribute *p*_*i*_ to additive variance would have polygenicity π_*i*_. (Strictly speaking, this would require π_*i*_ to be an integer, but we relax this requirement.) We denote the vector of variant polygenicities corresponding to a distribution *D* = (*p*_1_, …, *p*_*n*_) by Π_*D*_; namely, Π_*D*_ = (π_1_, …, π_*n*_) = (1⁄*p*_1_, …, 1⁄*p*_*n*_).

We would like a measure of polygenicity to be some kind of average of variant polygenicities. We would also like the contribution of a given variant to this average to be weighted by its proportional contribution to heritable variance, so that variants with nearly-zero contributions to variance do not dominate the average. We consider the three heritability-weighted Pythagorean means (arithmetic, harmonic, and geometric) as examples.

The *arithmetic mean* of variant polygenicities is

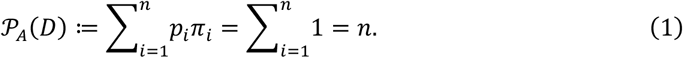

It equals the number variants with nonzero contributions to variance, which, as we already described, is often used as a measure of polygenicity.

The *harmonic mean* gives rise to the *effective polygenicity* proposed by O’Connor and colleagues^21^:

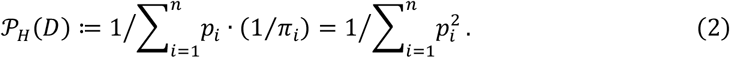

If *p* is the largest contribution to heritable variance then this measure is bound from below by 1⁄*p*^2^. O’Connor and colleagues showed that the prediction accuracy of the optimal polygenic score constructed from a GWAS with a small sample size is inversely proportional to the effective polygenicity^21^. This measure was called the ‘effective number of independently associated SNPs’ in ref. ^21^.

The *geometric mean* gives rise to a new measure, which we call the *entropy polygenicity*:

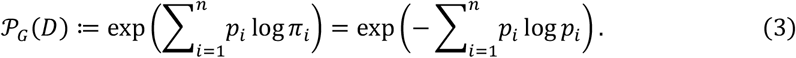

It does not depend on the base in which the log is taken. The exponent is formally equal to Shannon’s entropy in information theory and to entropy in statistics and statistical physics^22^. Akin to how entropy quantifies the diffusiveness of probability distributions in these fields, the entropy polygenicity quantifies the diffusiveness of the heritability distribution *D*.

### A general definition

The Pythagorean means are special cases of *quasi-arithmetic means*^23^. A quasi-arithmetic mean is defined in terms of a continuous and invertible function *f* between two intervals of the real numbers. The weighted quasi-arithmetic mean of *Z* = (*z*_1_, …, *z*_*n*_) with weights *W* = (*w*_1_, …, *w*_*n*_) such that 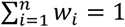 takes the form

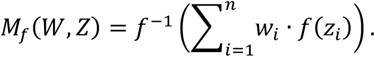

The Pythagorean polygenicity measures that we have considered are quasi-arithmetic means of the polygenicity of variants, Π_*D*_, weighted by their proportional contributions to heritable variance, *D*, i.e.,

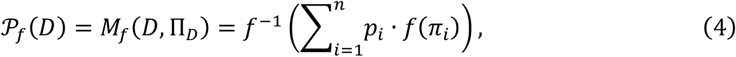

where *f*(*x*) = *x* for the arithmetic mean, *f*(*x*) = 1/*x* for the harmonic mean, and *f*(*x*) = log *x* for the geometric mean.

We will define polygenicity measures in terms of variant contributions to phenotypic rather than heritable variance. Namely, the *p*_*i*_’s will now denote variants’ absolute as opposed to relative contributions to phenotypic variance, and therefore they sum to the genetic variance, *V*_*A*_, rather than 1. If the phenotypic variance is one, then in these terms, the variant polygenicity, π_*i*_ = 1⁄*p*_*i*_, corresponds to the number of contributions of size *p*_*i*_ that would make up the phenotypic variance in the trait.

We define a polygenicity measure as one that takes the form

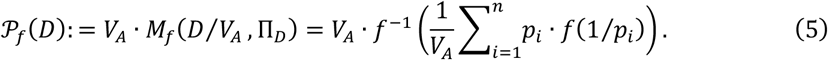

When we express measures in terms of individual contributions, we use the notation

*P* _*f*_(*D*) = *P* _*f*_(*p*_1_, …, *p*_*n*_). In this definition, *M*_*f*_(*D*⁄*V*_*A*_, Π_*D*_) is the quasi-arithmetic mean of variant polygenicities (defined relative to the phenotypic variance) weighted by their contribution to heritable variance. Multiplying the mean by *V*_*A*_ restores our original interpretation of polygenicity as the number of variants with the mean contribution that make up the total heritable variance.

For Pythagorean means this definition is equivalent to the one we used in Eq. (4), because these means satisfy the linear scaling property that *M*_*P*_(*W, λ · Z*) = *λ · M*_*P*_(*W, Z*) for any real *λ*, in particular *V*_*A*_. We do allow for polygenicity measures whose corresponding means do not satisfy this scaling property, and an example of such a measure is given in Appendix A.

### Defining properties

All polygenicity measures share mathematical properties, which derive from properties of quasi-arithmetic means^23^. Here, we list five properties and argue why they are sensible, perhaps even necessary, for a measure of polygenicity. In Appendix B, we show that any measure that satisfies these properties is a polygenicity measure by our definition. In other words, we can define a polygenicity measure by these properties.

The defining properties of polygenicity measures are:

1. ***Maximality*:** *P* (*p*_1_, …, *p*_*n*_) ≤ *n*, with equality when *p*_1_ = ⋯ = *p*_*n*_.
2. ***Symmetry*:** *P* (*p*_1_, …, *p*_*n*_) is invariant to permutations of its arguments (the *p*_*i*_’s).
3. ***Continuity*:** *P* (*p*_1_, …, *p*_*n*_) is a continuous function of its arguments (the *p*_*i*_’s).
4. ***Replacement*:** If *p*_1_ + ⋯ + *p*_*n*_ = *q*_1_ + ⋯ + *q*_*m*_ > 0 and *P* (*p*_1_, ⋯, *p*_*n*_) = *P* (*q*_1_, ⋯, *q*_*m*_), then for any *z* > 0, *P* (*z, p*_1_, ⋯, *p*_*n*_) = *P* (*z, q*_1_, ⋯, *q*_*m*_).
5. ***Convergence:*** If *p*_1_ = *p*_2_ = ⋯ = *p* > 0, then for any additional contribution 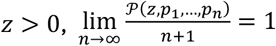.

The intuition behind the first three properties is straightforward. Maximality assures that when heritable variance is equally distributed among *n* variants the polygenicity is equal to *n*, and that this is the highest polygenicity attainable with *n* variants. Symmetry assures that the order in which variants are listed does not affect the polygenicity. Continuity assures that a small change in a variant’s contribution has a small effect on the polygenicity.

Because *P* is defined over sequences of positive numbers, the continuity property does allow for a discontinuity in the limit that variant contributions approach zero. This allows the arithmetic polygenicity—the number of nonzero contributions—to satisfy our definition, but it also gives rise to the statistical and conceptual problems with this measure (see Introduction).

The next two properties require more elaborate explanations but are important, nonetheless. Suppose that we divide a set of variants into two subsets, with contribution *p*_1_, …, *p*_*m*_ and *p*_*m*+1_, …, *p*_*n*_. We would expect the polygenicity of the entire set, *P* (*p*_1_, …, *p*_*n*_), to remain the same if we replace the first subset of variants with a different one with the same polygenicity and contribution to variance. This is the replacement property.

To better understand the convergence property, we consider an example. Suppose that heritable variance in a trait is equally distributed among *n* ‘small’ variants, and we add one ‘large’ variant. If a small variant contributes *p* to the absolute phenotypic variance and the large variant contributes *z* = 10*p*, then the large variant makes a proportional contribution *z*/(*z* + *np*) = 10/(10 + *n*). When *n* is sufficiently large, this contribution approaches zero, and we would expect the additional variant to have little effect on polygenicity. However, consider a measure that takes the form

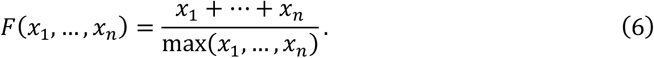

In this case, when we increase *n, F*(*z, p*_1_, …, *p*_*n*_) = *F*(10*p, p*, …, *p*) approaches *n*⁄10, rather than *n* + 1. This measure is extremely sensitive to the magnitude of the largest individual contribution, even if its proportional contribution to phenotypic variance is tiny. It is not a polygenicity measure by our definition: it satisfies properties 1-4 but not property 5. The convergence property precludes polygenicity measures from having such extreme sensitivity.

#### Softmax polygenicity

We might, nonetheless, be particularly interested in the largest contributions to variance. To this end, we use a continuous approximation of the max function in Eq. 6. We define the *softmax polygenicity* using *f*(*x*) = exp(−*x*), i.e.,

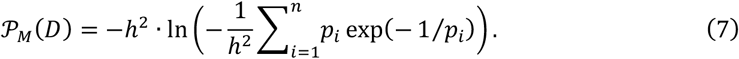

This measure approximates the variant polygenicity of the largest single contribution, provided that its proportional contribution is not negligibly small. For example, with *p*_1_ = 1/2, *p*_2_ = 1/4 and *p*_3_ = 1/4, the softmax polygenicity is − ln(*e*^−2^⁄2 + *e*^−4^⁄4 + *e*^−4^⁄2) ≈ −ln(*e*^−2^⁄2) ≈ 2.3. The softmax polygenicity has an approximate version of the linear scaling property: if max *D* ≫ 0 and *λ* > 1, then 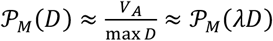.

### Entropy polygenicity of a partition

We often partition variants into subsets. Variants are partitioned into those that are coding and noncoding; coding variants are partitioned by their effect on the protein; noncoding variants are partitioned by their proximity to genes or predicted regulatory elements; additionally, variants are partitioned by their allele frequencies. Heritability is often partitioned across subsets of variants. Likewise, we would like our measure of polygenicity to apply to partitions in a natural and interpretable way.

To see how such an application would look like, consider a partition of *n* variants into *m* annotations. Such a partition defines three kinds of polygenicities, which we consider in turn. For simplicity, we assume a polygenicity measure that satisfies the linear scaling property and use the definition given in Eq. 4, with *p*_*i*_’s denoting proportional contributions to heritable variance; generalizing what follows using the definition given in Eq. 6 is straightforward (Appendix C).

We first consider the polygenicity of a single annotation *a* ∈ {1, …, *m*} with *n*_*a*_ variants. By analogy with the case with all variants affecting a trait, we denote the proportional contribution of variant *i* ∈ {1, …, *n*_*a*_} to the heritable variance explained by this annotation by 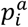, and the distribution of variance among variants in this annotation by 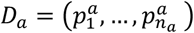, where 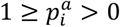 and 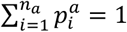. We therefore define the polygenicitywithin this annotation as

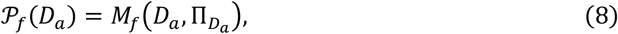

with *f* defining the specific measure.

Next, we consider the mean polygenicity over the annotations in the partition. We denote the proportional contribution of annotation *a* ∈ {1, …, *m*} to heritable variance—the sum of contributions over the *n*_*a*_ variants in it—by *p*^*a*^; the distribution of these contributions by

*D*_*A*_ = (*p*^1^, …, *p*^*m*^), where 1 ≥ *p*^*a*^ > 0 and 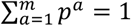 and the vector of polygenicities associated with these annotations by Π_*A*_ = (*P*_*f*_(*D*_1_), …, *P*_*f*_(*D*_*m*_)), with *P*_*f*_(*D*_*a*_) for *a* = 1, …, *m* defined in Eq. 8. Following our original reasoning with variants, we take the quasi-arithmetic mean over the polygenicity of annotations weighted by their proportional contributions to heritable variance; namely,

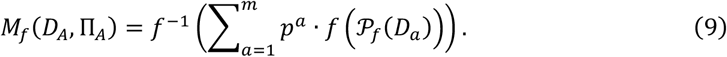

Lastly, we consider the polygenicity of the partition; namely, the polygenicity associated with the distribution of heritable variance among annotations, *D*_*A*_ = (*p*^1^, …, *p*^*m*^). Again, by analogy, we define an annotation’s polygenicity as π^*a*^ ≡ 1⁄*p*^*a*^, and denote the vector of annotation polygenicities by 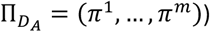. We define the polygenicity of the partition as

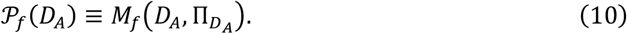

We would like the polygenicity of the total set of variants to equal the polygenicity of the partition times the mean polygenicity of the annotations in it. Namely, we would like our measure of polygenicity to satisfy

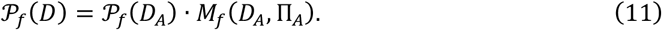

In appendix C, we show that the entropy polygenicity satisfies this requirement; we also conjecture that it is the only polygenicity measure that does.

### Linkage disequilibrium

Linkage disequilibrium (LD) affects how phenotypic variation is distributed in the population and should therefore affect measures of polygenicity. Consider, for example, two variants in perfect LD, with alleles a and A in one and b and B in the other, where only haplotypes ab and AB are present in the population. The effect of these variants on phenotypic variation is equivalent to the effect of a single variant with alleles ‘ab’ and ‘AB’. We would therefore like for the joint contribution of the two variants to polygenicity to equal the contribution of such an effective variant. Moreover, when the variants are in partial LD, we would like their contribution to polygenicity to lie in between the contributions in the extremes of perfect LD and LE.

In Appendix D, we generalize our definition of a polygenicity measure to account for LD. We do so in the context of a random effects model, following ref. ^21^. We distinguish between direct, or causal, contributions to variance (*p*_*i*_), and marginal contributions, *z*_*i*_. If *p*_*i*_ is the variance of a causal effect size and *z*_*i*_ is the variance of a marginal effect size, then these are related by the formula 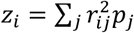, with LD matrix 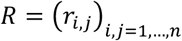. We use the marginal contributions to define the variant polygenicities, Π_*Z*_ = (1⁄*z*_1_, …, 1⁄*z*_*n*_), and the direct contributions, *D* = (*p*_1_, …, *p*_*n*_), to define their weights. Namely, a polygenicity measure is now a function of the causal contributions and the LD matrix, and it takes the form

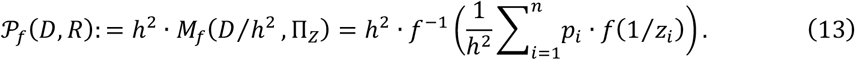

This definition aligns with our intuitions about the effect of LD on polygenicity (see Appendix D). When all variants are in LE it reduces to our previous definition, because *z*_*i*_ = *p*_*i*_ for any variant *i*. In the example of two variants in perfect LD, both variants (A/a and B/b) have the same variant polygenicity, which accounts for the sum of their effect sizes, and their weights sum up to their (expected) combined contribution to variance. Their combined contribution to polygenicity equals the (expected) contribution of a single effective variant (AB/ab). More generally, we show that this definition aligns with our intuition when variants are arranged in perfect LD blocks (roughly approximating human LD structure).

### Estimating polygenicity

We use Fourier Mixture Regression (FMR)^24^ to estimate polygenicity from GWAS summary statistics. FMR estimates the fraction of common variant heritability that is explained by variants with different marginal effect sizes. It fits a mixture of mean-zero Normal distributions *k* = 1, …, *K*, with weights *w*_*k*_ that are estimated from the data and variances 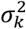 that are fixed. *w*_*k*_ quantifies the fraction of heritability explained by mixture component *k*. FMR estimates the weights by matching the empirical characteristic function of the GWAS z-scores—the Fourier transform of their density function—with its expectation under the mixture model. We show that a polygenicity measure (defined by a function *f*) is estimated by

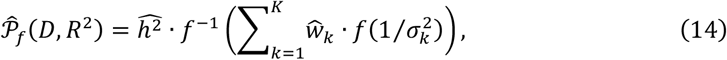

where 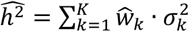. Details on the estimate and estimation are provided in Appendix E.

## Results

### Relationship among polygenicity measures

The four polygenicity we analyze measures satisfy this relationship, which can be derived using the AM-GM inequality:

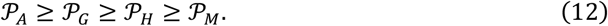

For a trait whose causal variants all contribute equally to variance, the four measures are equal. To illustrate how the four measures differ, therefore, we consider a trait whose contributions to variance are either ‘small’, with size *p*_*s*_, or ‘large’, with size *p*_*l*_ ≥ *p*_*s*_. We vary *p*_*s*_, *p*_*l*_, and the number of contributions, *n*_*s*_ and *n*_*l*_.

In Fig. 1A we increase *n*_*s*_ and decrease *p*_*s*_ such that the total contribution *n*_*s*_*p*_*s*_ remains constant. As *n*_*s*_ increases, *P* _*A*_ and *P* _*G*_ increase without bound, whereas *P* _*H*_ and *P* _*M*_ are bounded. In such cases, *P* _*G*_ ≫ *P* _*H*_. This example resembles the one we considered in the Introduction, except that the sum of the small effects is non-negligible; if the sum of the small effects went to zero instead of remaining constant, then *P* _*G*_ would be bounded.

**Figure 1.**
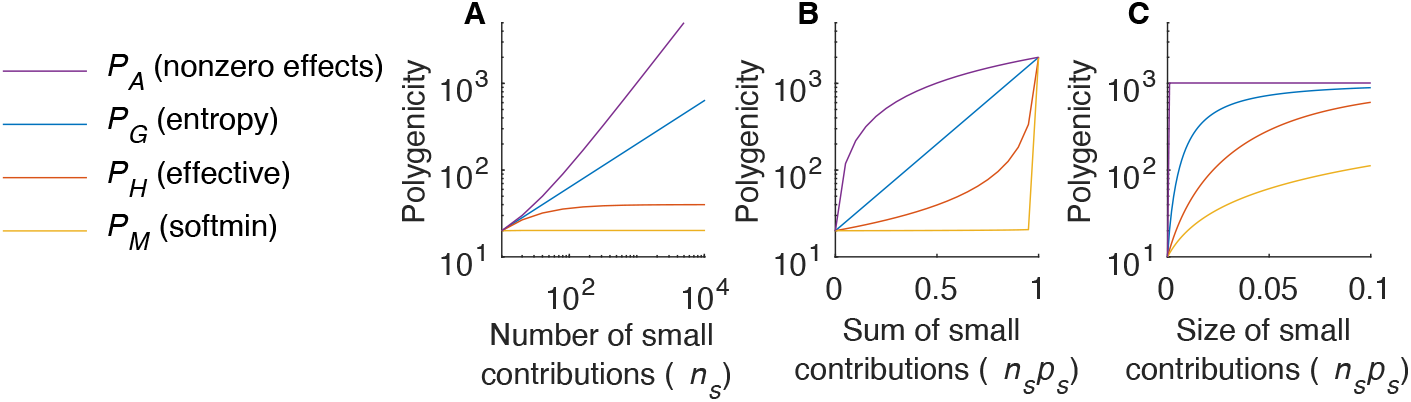
Comparison of polygenicity measures. The four measures are calculated across a range of simulated genetic architectures involving a mixture of small and large contributions. In (A) the number of small contributions varies while their sum remains constant. In (B) the sum of small contributions varies with the number of them, while their sizes remain constant. In (C) the sum of small contributions varies with their sizes, while the number of them remains constant. In (A) we fix *n*_*l*_ = 10, *p*_*l*_ = 0.01 and *n*_*l*_*p*_*l*_ = *n*_*s*_*p*_*s*_ = 0.1, and vary *n*_*s*_ and thus *p*_*s*_. In (B) we fix *p*_*s*_ = 10^−4^, *p*_*l*_ = 10^−2^ and *n*_*l*_*p*_*l*_ + *n*_*s*_*p*_*s*_ = 0.2, and vary *n*_*s*_ and thus *n*_*s*_*p*_*s*_⁄(*n*_*l*_*p*_*l*_ + *n*_*s*_*p*_*s*_). In (C) we fix *n*_*l*_ = 10, *p*_*l*_ = 0.01 and *n*_*s*_ = 10^3^ and vary *p*_*s*_⁄*p*_*l*_ and thus *n*_*s*_*p*_*s*_⁄(*n*_*l*_*p*_*l*_ + *n*_*s*_*p*_*s*_).

In Fig. 1B we increase *n*_*s*_ and decrease *n*_*l*_ while fixing the individual contributions, *p*_*s*_ and *p*_*l*_, and the total variance, *n*_*s*_*p*_*s*_ + *n*_*l*_*p*_*l*_. Here, the proportional contribution of small variants, *n*_*s*_*p*_*s*_, increases with *n*_*s*_. *P* _*A*_, *P* _*G*_ and *P* _*H*_ increase with the proportional contribution of small variants. In contrast, *P* _*M*_ remains nearly constant until there are no large effects remaining. Thus, when large contributions exist but contribute a small proportional of the variance, *P* _*H*_ ≫ *P* _*M*_.

In Fig. 1C we increase the individual contributions of small variants, *p*_*s*_, while fixing *n*_*s*_, *p*_*l*_, and *n*_*l*_. So long as *p*_*s*_ > 0, even if the total contribution of small variants is negligibly small, the total number of variants and thus *P* _*A*_ remain constant. In contrast, the other measures increase continuously with *p*_*s*_, converging to *P* _*A*_ when *p*_*s*_ = *p*_*l*_. Thus, when many small variants contribute negligibly to the total variance, *P* _*A*_ ≫ *P* _*G*_.

These examples show that there can be arbitrarily large differences between any two of the four polygenicity measures that we consider, motivating us to quantify more than one of them.

### Simulations

Three of our considered polygenicity measures can be quantified using an existing method, Fourier Mixture Regression^24^ (see Methods). To test this approach, we simulated summary statistics from their asymptotic sampling distribution (see Appendix F) and applied FMR to estimate their entropy polygenicity (*P* _*G*_), their effective polygenicity (*P* _*H*_), and their softmax polygenicity (*P* _*M*_). Estimates of *P* _*G*_ were slightly upwardly biased (Figure 2A), estimates of *P* _*H*_ were slightly downwardly biased (Figure 2B), and estimates of *P* _*M*_ were approximately unbiased (Figure 2C). These results, including the existence of bias, align with expectations; FMR should produce estimates that are consistent but not unbiased at finite sample size. For all three metrics, FMR estimates were consistently <0.5 orders of magnitude different from true values. Similar results were obtained in simulations where we varied the sample size, the simulated effect-size distribution, and the distribution of heritability across allele frequencies (Appendix F).

**Figure 2.**
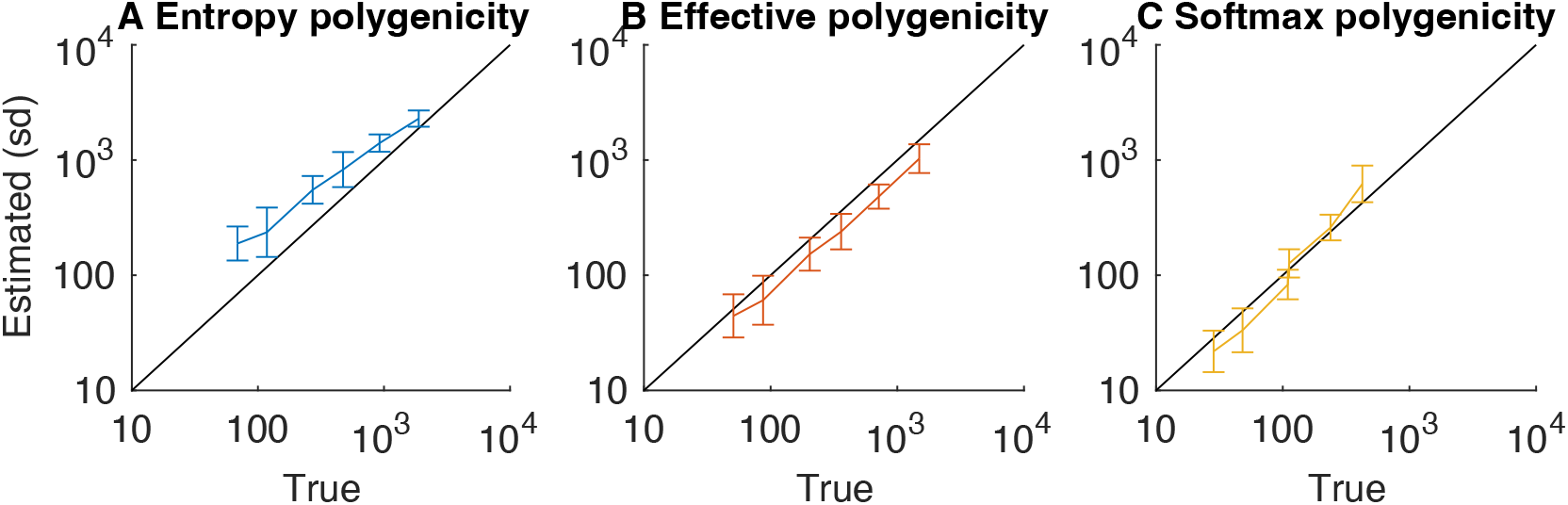
Polygenicity estimates vs ground truth in simulations. Mean and SE were estimated using 100 replicates of simulations with a given set of parameters. See Appendix F for details and Figure F1 for additional simulation results.

### Polygenicity of 36 traits in humans

We estimated the polygenicity of 25 quantitative traits and 11 diseases in individuals of European ancestry by applying FMR to the GWAS summary statistics of variants with MAF>1%. The traits we included span the major phenotypic categories commonly studies in GWAS. The quantitative traits include anthropometric traits (e.g., height and body mass index), cognitive/behavioral traits (e.g., years of education), quantitative medical traits (e.g., blood pressure), biomarkers and hematopoietic traits, a reproductive trait, and a pigmentation trait. The diseases include autoimmune and inflammatory diseases (e.g., inflammatory bowel disease), psychiatric diseases (e.g., schizophrenia), and coronary artery disease. The list of traits and their basic characteristics are detailed in Table S1.

For each trait, we estimated the entropy polygenicity (*P* _*G*_), the effective polygenicity (*P* _*H*_), and the softmax polygenicity (*P* _*M*_). As we would expect (Eq. 12), the estimates for all traits satisfy 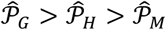. For most traits, 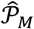 ranges between 50 and 500; 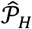 ranges between 500 and 10,000; and 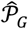 ranges between 5,000 and 100,000. The difference between 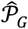 and 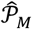 (approximately two orders of magnitude) indicates that much of the heritable variance in traits arises from variants whose individual contributions are much smaller than the largest ones. The difference between 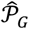 and 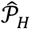 (one order of magnitude) suggests that heritable variance accumulates across a wide range of contribution sizes. Both findings are consistent with previous estimates of the effect size distribution ^24,25^.

Recent studies further suggest that the distributions of common variant selection coefficients and contributions to heritability are fairly similar among complex traits^24,25^. On the other hand, complex traits vary widely in their mutational target size^25^; if other factors are similar across traits, the mutation target size determines their polygenicity, and different polygenicity metrics are proportional to the target size, and to each other. Indeed, the three metrics are highly correlated (mean *r*^2^ = 0.78) (Fig. 3B-C), and they are correlated with estimates of the mutational target from Simons and colleagues^25^ (mean

**Figure 3.**
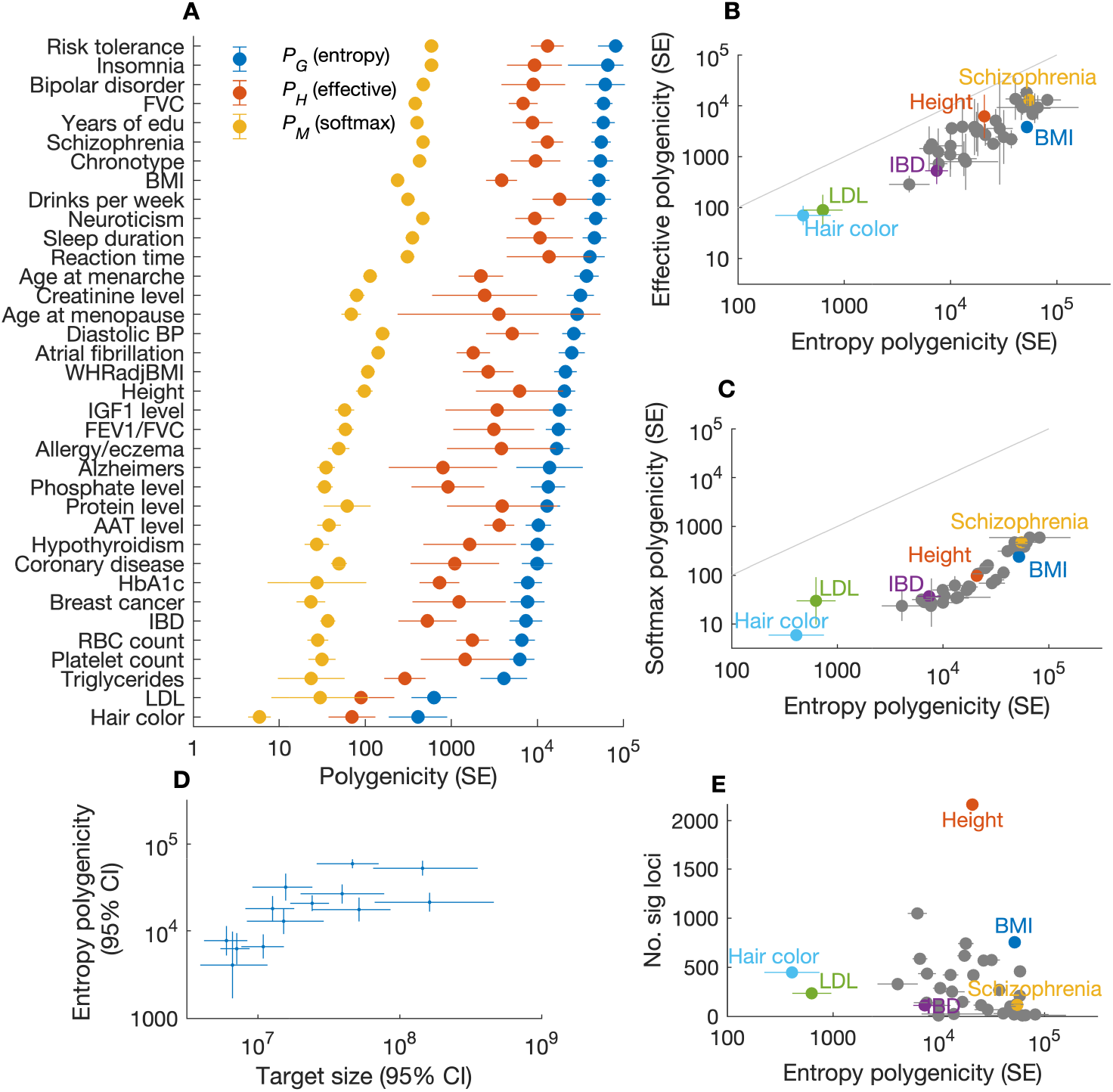
Estimates of polygenicity for 36 complex traits in humans. (A) Estimates *P* _*G*_, *P* _*H*_ and *P* _*M*_ across traits, sorted by 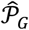. (B) Effective vs entropy polygenicity estimates. Line indicates y=x. (C) Softmax vs. entropy polygenicity estimates. Line indicates y=x. (D) Estimates of entropy polygenicity vs. estimates of mutational target size from ref. ^25^ for 13 quantitative traits, with 95% confidence intervals. (E) The number of genome-wide significant loci

*r*^2^ = 0.52) (Fig. 3D). These correlations are not adjusted for attenuation due to sampling variation. In contrast, the number of genome-wide significant loci for a trait strongly depends on the trait’s heritability per site, which varies considerably among traits^24^, and there is no apparent correlation between polygenicity and the number of GWAS hits (mean *r*^2^ = 0.02) (Fig. 3E).

Most of the traits that we analyzed can be called ‘highly polygenic’, with entropy polygenicity between 5,000 and 100,000. Only two traits, hair color and LDL cholesterol, had lower values of 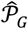, between 100 and 1,000 (Fig. 3A). For comparison, in Appendix G we approximate that a typical simple (Mendelian) trait would have *P* _*G*_ ≈ 4. This number is small because the trait polygenicity associated with a gene is on the order of *L · θ*, where *L* is the mutational target size of the gene associated with changes to the trait, and *θ* is the neutral diversity level; in humans, *L · θ* is on the order of 1 (Appendix G). Even allowing that apparently monogenic traits can have more complex inheritance, there appears to be a dearth of traits whose polygenicity is intermediate between simple traits, with *P* _*G*_ ≈ 4, and highly polygenic traits, with *P* _*G*_ > 5,000.

## Discussion

We proposed a single definition of polygenicity that encompasses many specific polygenicity measures. These different measures have different interpretations, making it useful to quantify multiple of them. For example, the effective polygenicity has an interpretation related to the prediction accuracy of polygenic scores^21^; differences in the effective polygenicity across traits may partially explain differences in the performance of risk scores. The softmax polygenicity is interpreted as the number of contributions similar to the largest single contribution that would be required to explain trait heritability; this makes it a natural choice of polygenicity metric when investigating top GWAS loci. In contrast, the entropy polygenicity reflects an average over the full range of contribution sizes, not emphasizing any particular scale, and it has the advantage that it applies to partitions in a natural and interpretable way. All of these metrics satisfy the properties that we consider to be necessary for a measure of ‘polygenicity’.

Our polygenicity estimates included the contributions of common variants only. Common variants likely explain most heritable variation in complex traits ^7,26^, and all of our polygenicity metrics are heritability-weighted averages, so the polygenicity of rare and common variant contributions combined is expected to be similar to that of common variants alone. On the other hand, rare variants on their own could have dramatically different polygenicity. O’Connor and colleagues found that the effective polygenicity is smaller for low frequency versus common variants^21^, and Weiner & Nadig and colleagues found that when rare coding variants are aggregated by gene, individual genes often explain >1% of the total variance explained, consistent with rare variants having much lower polygenicity^26^.

We find a dearth of traits with intermediate polygenicities, and this observation seemingly requires explanation. Plausibly, it could reflect the ascertainment of traits in human GWAS, which is often driven by biomedical and other biological considerations. There are considerable redundancies among traits, including for the traits we examined, and many traits are not represented either in our study or in GWAS more generally. However, trait ascertainment bias might be expected to cause overrepresentation, rather than the opposite, of traits with intermediate polygenicity. GWAS would be especially productive for such traits because they have many loci to discover, yet those loci are detectable at modest sample sizes. If GWAS are biased toward the traits for which they are most productive, it would not explain the apparent dearth of intermediate polygenicities that we observe. A different explanation is that in humans, Mendelian diseases are often syndromic, producing a recognizable constellation of symptoms that may ‘fingerprint’ a single causative gene or pathway. If polygenicity were quantified for symptoms as opposed to syndromes, and each symptom could be caused by numerous syndromes, then the polygenicity gap might shrink. For example, intellectual disability is a common feature of many developmental disorders, and over 1,000 genes are thought be involved in early onset developmental disorders in aggregate^27^.

More generally, our findings highlight two questions that have received considerable recent attention: (i) why are the traits considered in human GWASs as highly polygenic as they are, and (ii) what determines variation in polygenicity among them. Part of the answer is plausibly evolutionary. Under certain relationships between variant effects on traits and on fitness (and everything else, notably pleiotropic effects, being equal), variant sites with effect sizes exceeding a tiny value are expected to have similar contributions to trait heritability^28,29^ (see also refs. ^30,31^). If there exist far more sites with small versus large effect sizes, this threshold effect causes increased polygenicity, a phenomenon termed ‘flattening’^21^. Flattening would cause polygenicity to be greater for common variants versus rare and low-frequency variants, consistent with what is observed^21,26^, and models of stabilizing selection that predict flattening have been shown to fit the data from human GWASs very well^25,28,32^. Flattening helps to explain why a trait with a large mutational target should have high polygenicity despite also having a handful of sites with much larger effect sizes. However, it does not explain why the mutational target should be so large to begin with.

The omnigenic model suggests a tentative explanation^16,33,34^. Consider a trait whose value is directly affected by the activity of a modest number of ‘core’ genes in particular cellular contexts (e.g., cell types and life history stages). The omnigenic model posits (and evidence is supportive) that many of the variants that directly affect the activity of any gene in the relevant cellular contexts would affect the activity of the core genes and thus contribute to heritable variation in the trait. The model explains the high polygenicity of traits by there being many such variants. It could explain variation in polygenicity among traits as arising from variation in the numbers of core genes and active regulatory regions and ‘peripheral’ genes in the cellular contexts that affect traits, as well as from variation in the number of cellular contexts that affect them. For example, morphological and brain related traits might be substantially more polygenic than molecular traits (including LDL, which was one of our two outliers) partially due to the greater number of cellular contexts that affect them. Additionally, one could imagine that the activity of core genes of some traits could be better insulated from their cellular contexts than those of other traits (e.g., via modularity and regulatory feedback), and are therefore affected by fewer regulatory variants (that exceed the flattening threshold).

The combination of evolutionary flattening and omnigenic regulatory dynamics provides a plausible framework for thinking about differences in polygenicity among complex traits. Nonetheless, we still have ways to go to understand the genetic underpinning of traits and how those relate to heritable variation in them. Other mechanisms might also be at play. For example, a long-term balance between local adaptation and migration could generate genetic architectures in which few major loci contribute a considerable fraction of heritable variance^35–39^. It has been speculated that such migration-selection balance plays a central role in other species and might have contributed to the low polygenicity of pigmentation traits is humans (possibly including hair color, which was our most extreme outlier)^35,39,40^. Being able to estimate well-defined measures of polygenicity for different kinds of traits should be helpful in better understanding what determines trait polygenicity.

## Acknowledgements

We thank Michael Lachman for discussions about the relationship between entropy polygenicity and Shannon’s information. This work was supported by NIH-NIGMS grant R35GM155278 to LJO and NIH R01 grant GM115889 to GS.

## Code availability

Fourier Mixture Regression software is available at https://github.com/lukejoconnor/FMR. Software to simulate GWAS summary statistics using realistic LD patterns is available at https://github.com/oclb/graphld.

## Data availability

GWAS summary statistics are available at https://console.cloud.google.com/storage/browser/broad-alkesgroup-public-requester-pays;tab=objects?prefix=CforceOnObjectsSortingFiltering=false.

## Appendix A threshold polygenicity

We may be interested in some kind of “threshold polygenicity”: the number of variants whose contributions to phenotypic variance exceed a threshold *t* ≪ 1. This would be a sensible alternative to ‘the number of nonzero effects’, potentially overcoming the problem that it is sensitive to arbitrarily small contributions (see Introduction). For the new measure to be continuous at the threshold, we can ‘smoothen’ its dependance on variant contributions near the threshold using an invertible activation function. For example, let

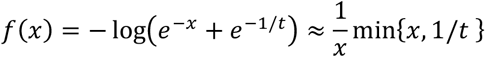

for which *xf*(*x*) ≈ 1/*t* for variants with effective polygenicity *x* ≳ 1/*t*, and *xf*(*x*) ≈ *x* for variants with effective polygenicity *x* ≪ 1/*t*. If all non-null variants have contributions *p* greater than *t* (effective polygenicities less than 1/*t*), then *f* is approximately the identity function, and its corresponding polygenicity measure approximates the number of nonzero effects, *P* _*A*_. Contributions smaller than the threshold are counted as a fraction of a contribution: a contribution of size *t*/2 is counted as approximately one half of a contribution, and so on.

The threshold polygenicity does not have the linear scaling property, as it depends on the size of the contributions *p*_1_, …, *p*_*n*_ not in relation to each other but rather in relation to *t*.

## Appendix B defining properties of polygenicity measures

Let *M* be a real-valued function on the finite sequences of real numbers, *x*_1_, …, *x*_*n*_. *M* is called a *quasi-arithmetic mean* if it has the following properties:

i. *M*(*x*, …, *x*) = *x*
ii. *M* is permutation invariant in its arguments
iii. *M* is continuous in each of its arguments
iv. If *M*(*x*_1_, …, *x*_*n*_) = *M*(*y*_1_, …, *y*_*n*_), then *M*(*x*_1_, …, *x*_*n*_, *z*) = *M*(*y*_1_, …, *y*_*n*_, *z*)
v. *M* is monotonically increasing in each of its arguments.

Kolmogorov^23^ showed the following:

**Theorem 1 (characterization of quasi-arithmetic means)**

*If M is a quasi-arithmetic mean, then there exists a continuous, invertible function f*: ℝ → (0, ∞) *such that:*

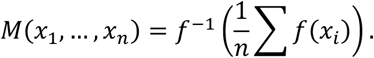

We show that quasi-arithmetic means are related to polygenicity functions, as follows. Let *P* be a real-valued function on the finite sequences of positive numbers, *x*_1_, …, *x*_*n*_, with *n* > 0. We call *P* a *polygenicity function* if it has the following properties:

1. Maximality: *P* (*x*_1_, …, *x*_*n*_) ≤ *n*, with equality when *x*_1_ = *x*_2_ = ⋯ = *x*_*n*_.
2. Symmetry: *P* is permutation invariant in its arguments
3. Continuity: *P* is continuous in its arguments
4. Replacement: If *x*_1_ + ⋯ + *x*_*n*_ = *y*_1_ + ⋯ + *y*_*m*_ and *P* (*x*_1_, …, *x*_*n*_) = *P* (*y*_1_, …, *y*_*m*_) then *P* (*x*_1_, …, *x*_*n*_, *z*) = *P* (*y*_1_, …, *y*_*n*_, *z*)
5. Convergence: if *x*_1_ = *x*_2_ = ⋯, then for any *y* > 0:

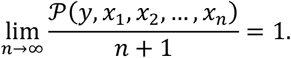

Properties 1-3 of a polygenicity function coincide one-to-one with those of a quasi-arithmetic mean. The relationship between properties 4-5 is not quite so apparent, but nonetheless, we show the following:

**Theorem 2 (characterization of polygenicity functions)**

*If P is a polygenicity function, then there exists a continuous, monotonically increasing function f*: (0, ∞) → (0, ∞) *such that:*

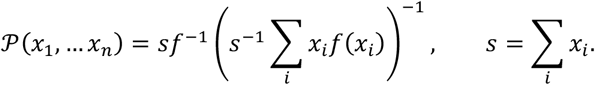

**Proof**. Let *P* be a polygenicity function, and let:

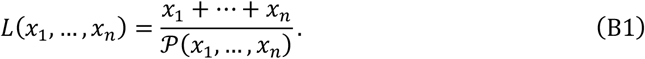

*L* has the following 5 properties, inherited from properties 1-5 of a polygenicity function respectively:

1. 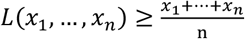, with equality when *x*_1_ = *x*_2_ = ⋯ = *x*_*n*_.
2. *L* is permutation invariant
3. *L* is continuous in each of its arguments
4. If *x*_1_ + ⋯ + *x*_*n*_ = *y*_1_ + ⋯ + *y*_*m*_ and *L*(*x*_1_, …, *x*_*n*_) = *L*(*y*_1_, …, *y*_*m*_) then *L*(*x*_1_, …, *x*_*n*_, *z*) = *L*(*y*_1_, …, *y*_*n*_, *z*).
5. If *x*_1_ = *x*_2_ = ⋯ = *x*, then for any *y* > 0:

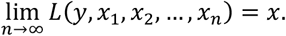

However, property (4) makes it clear that *L* is a weighted mean, with weights equal to the objects being averaged. We convert *L* into a function *M* that will be shown to be a quasi-arithmetic mean. We begin by defining *M*(*x*_1_, *x*_2_) for two integers. For *x*_1_, *x*_2_ ∈ ℤ, let:

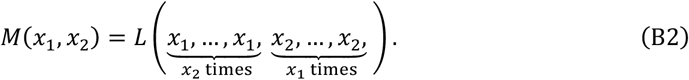

This definition generalizes to the mean of several integers *x*_1_, …, *x*_*k*_ by repeating *x*_*k*_ *a*_*k*_ = ∏_*j* ≠ *k*_ *x*_*j*_ times. It generalizes to the mean of several rational numbers by multiplying *a*_1_, …, *a*_*n*_ through by the product of their denominators, *d*, such that *x*_*k*_ is repeated *d* ∏_*j* ≠ *k*_ *x*_*j*_ times. This is acceptable because if the entire sequence is repeated, it has no effect: let *m* = *L*(*x*_1_, …, *x*_*n*_). Then by properties 1 and 4, *L*(*x*_1_, …, *x*_*n*_, *x*_1_, …, *x*_*n*_) = *L*(*m*, …, *m, m*, …, *m*) = *m*.

Having defined *M* over ℚ, we show that it has the replacement property (iv). Let *x*_1_, *x*_2_, *y*_1_, *y*_2_, *z* ∈ ℚ, and suppose that *M*(*x*_1_, *x*_2_) = *M*(*y*_1_, *y*_2_). Let *d* be the product of their denominators, and let *a* = *x*_1_*x*_2_*y*_1_*y*_2_*zd* ∈ ℤ. Then:

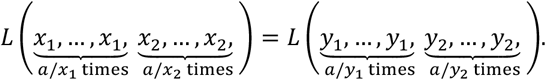

Therefore,

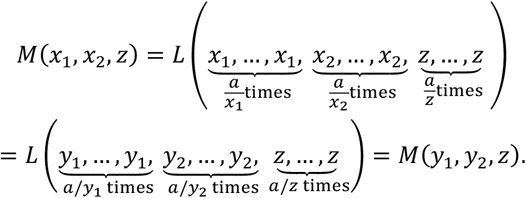

because the sequences being substituted have the same *L*-mean and the same sum (which is equal to 2*a*). This shows that *M*(*x*_1_, *x*_2_, *z*) = *M*(*y*_1_, *y*_2_, *z*).

Next, we show that if *y* > *x* ∈ ℚ, then:

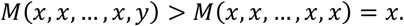

The property that *L*(*x*_1_, …, *x*_*n*_) ≥ ^*x*1+⋯+*xn/n*^ implies that:

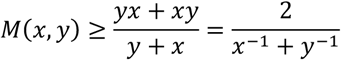

and more generally that:

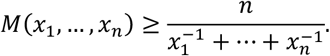

That is, whereas *L* is at least the arithmetic mean, *M* is at least the harmonic mean. We find that if *y* > *x*, then:

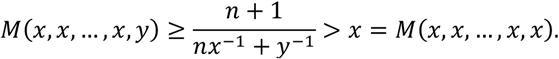

This property allows us to define *M* for a pair of real numbers. Consider the mean *M*(*x, y*) where *x* is an integer, *y* is irrational, and *x* > *y*. Let 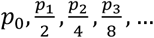 be a sequence of rational approximations to *y* such that for all *i*,

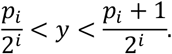

Let:

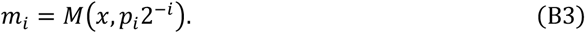

*m*_1_, *m*_2_, … is an increasing sequence, because

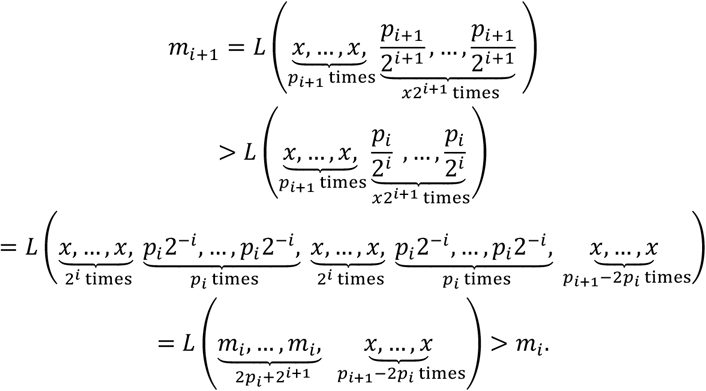

Moreover, *m*_1_, *m*_2_, …, is bounded, so the sequence converges. We define:

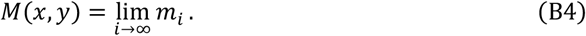

This definition extents to pair of irrational numbers (or any number of them) by taking a sequence of rational approximations to each irrational argument in turn.

This definition, together with the continuity of *L*, shows that *M* is continuous. It is also permutation invariant and idempotent. We already showed that it has the subsequence replacement property on ℚ, which extends to the reals by continuity. Moreover, we already showed that it is monotonically increasing at a point *M*(*x, x*). It remains to show that it is monotonically increasing elsewhere.

First, we need to show that for any *y, z* ∈ (0, ∞), there exists *x*_1_, …, *x*_*n*_ such that

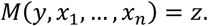

Suppose that *y* < *z*, and choose *x* > *z*. Let *x*_1_ = *x*_2_ = ⋯ = *x*. By the convergence property, we know that for some *n, M*(*y, x*_1_, …, *x*_*n*_) > *z*. Due to continuity, and by the intermediate value theorem, there exists *x*^′^ ∈ (*z, x*) such that *M*(*y, x*′, …, *x*′) = *z*.

Finally, we show that *M*(*y, z*) is an injective function of *y*. Suppose that *M*(*y, z*) = *M*(*y*^′^, *z*). There exists *x*_1_, …, *x*_*n*_ such that *M*(*z, x*_1_, …, *x*_*n*_) = *y*. Let *m* = *M*(*y*^′^, *y, y*, …, *y*). By subsequence replacement:

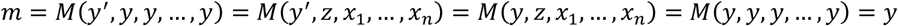

*m* = *M*(*y*^′^, *y, y*, …, *y*) = *M*(*y*^′^, *z, x*_1_, …, *x*_*n*_) = *M*(*y, z, x*_1_, …, *x*_*n*_) = *M*(*y, y, y*, …, *y*) = *y* As we already showed, this implies *y*^′^ = *y*, and *M* is an injection. A continuous injection is monotonic, and *M* is increasing at the point (*y, y*), so it is monotonically increasing.

This shows that for any polygenicity function *P*, there exists a corresponding quasi-arithmetic mean *M*. By Theorem 1, there exists a continuous, monotonically increasing function *f*: (0, ∞) → (0,1) such that:

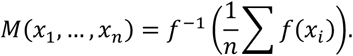

Working backwards, we find that in the rational case, if *x*_1_, …, *x*_*n*_ have common denominator *d*,

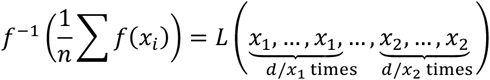

such that *L* can be written:

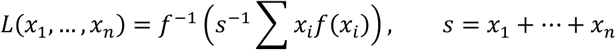

and *P* = *s*/*L* can be written:

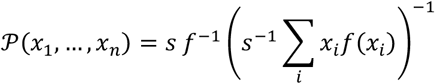

as desired.

## Appendix C Entropy polygenicity of a partition

Here we show that the entropy polygenicity satisfies the desired requirement on partitions (Eq. 11); namely, that

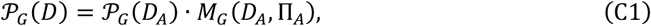

where:

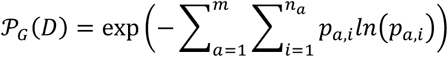

is the total polygenicity, with *p*_*a,i*_ 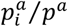 being the contribution of variant *i* in annotation *a* to heritable variance in the trait;

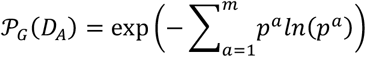

is the polygenicity of the partition, with 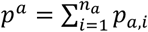 being the contribution of annotation *a* heritable variance in the trait;

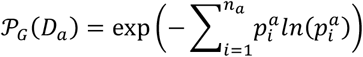

is the polygenicity of annotation *a*, with 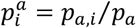 being the contribution of variant *i* in annotation *a* to heritable variance in this annotation; and

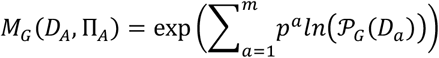

is the mean polygenicity over the annotations in the partition, with Π_*A*_ = (*P* _*G*_(*D*_1_), …, *P* _*G*_(*D*_*m*_)). To show that the Entropy polygenicity satisfies the requirement on partitions, we start from the mean polygenicity over annotations:

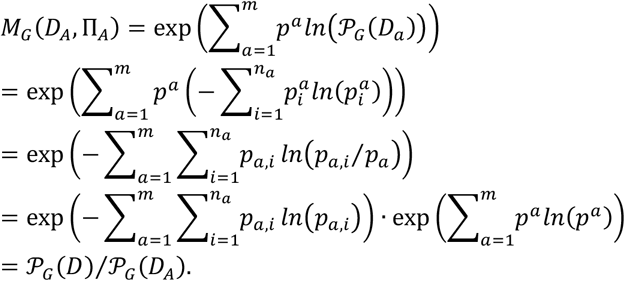

Using our general definition of polygenicity—without assuming that the mean under consideration satisfies the scaling property—the requirement on partitions takes the form

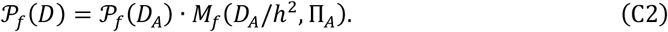

In the entropy polygenicity case, the terms take the following form:

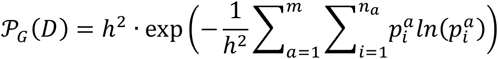

is the total polygenicity, with 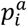 being the contribution of variant *i* in annotation *a* to phenotypic variance in the trait, and *h*^2^ being the heritability of the trait;

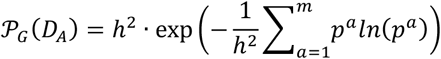

is the polygenicity of the partition, with 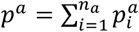 being the contribution of annotation *a* phenotypic variance in the trait;

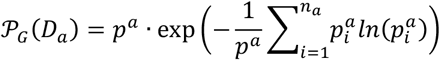

is the polygenicity of annotation *a*, with *p*^*a*^⁄*p*^*a*^ being the contribution of variant *i* in annotation *a* to heritable variance in this annotation; and

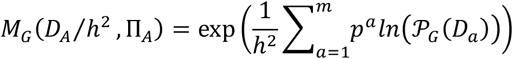

is the mean polygenicity over the annotations in the partition, with Π_*A*_ = (*P* _*G*_(*D*_1_), …, *P* _*G*_(*D*_*m*_)). Similar our previous derivation, we show that the Entropy polygenicity satisfies the requirement on partitions by starting from the mean polygenicity over annotations:

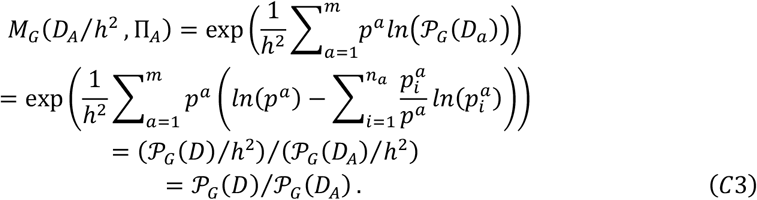

Our requirement on partitions resembles the grouping requirement in information theory (see, e.g., pp. 49-50 in ref. ^22^). The grouping requirement along with requirements that are analogous to our defining properties 1 and 3 (in main text) have been proved to uniquely define Shannon’s entropy (see, e.g., p. 54 in ref. ^22^; also see ref. ^41^). This is why we conjecture that the entropy polygenicity is the only polygenicity measure that satisfies our requirement on partitions.

## APPENDIX D definition of polygenicity with LD

We define polygenicity measures with LD in the context of a random effects model, the non-i.i.d. normal model. We assume that the vector of phenotypes *y*, with columns corresponding to individuals, is given by

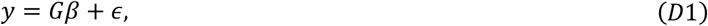

where *ϵ* is a vector of random environmental contributions uncorrelated from *Gβ*, and we define the genotype matrix *G* and vector of effect sizes *β* as follows. As before, we assume that causal variants are biallelic. We use the standard definition of the genotype matrix *G*, with rows corresponding to individuals, columns corresponding to causal loci, and entries being the standardizes genotypes of an individual at a locus. Namely, given MAF *f*_*j*_ at locus *j*, the standardized genotypes at the locus are 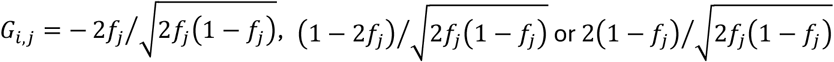. We measure the phenotype in units of the phenotypic standard deviation 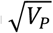. In these units and with standardized genotypes, the effect size at locus *j* takes the form 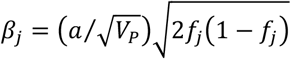, where *β*^*T*^ = (*β*_1_, …, *β*_*n*_). In this formulation, the random effects assumption takes the form

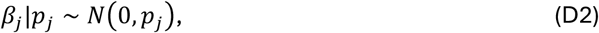

with *j* = 1, …, *n*. While we assume that effects are random rather than fixed, the model makes minimal assumptions about the genetic architecture. We use it because it simplifies the mathematical treatment and estimation that follow.

We define polygenicity measures in terms of variants’ direct and marginal contributions to phenotypic variance. The expected, direct, proportional contribution to variance of variant *j*—in the absence of LD—is *p*_*j*_. The expected variance contributed by all variants—with 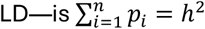; we define variant *j*’s direct contribution to heritability as *p*_*j*_ = *x*_*j*_⁄*h*^2^. With LD matrix *R* = (*r*)_*i,j*=1,…,*n*_, the expected marginal contribution to phenotypic variance of causal variant *j* is

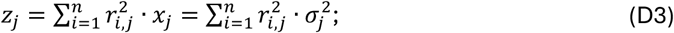

for brevity, we refer *z*_*j*_ as the marginal contribution. The realized marginal effect size 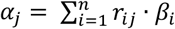 is normally distributed with *α*_*j*_∼*N*(0, *z*_*j*_).

Our definition of polygenicity measures uses direct contributions to heritability for weights and marginal contributions to phenotypic variance for the effective numbers of variants.

The weights are given by the vector *D*⁄*h*^2^ = (*p*_1_, …, *p*_*n*_) of direct contributions, whereas the effective numbers are based on the vector *Z* = (*z*_1_, …, *z*_*n*_) of marginal contributions, and are given by the vector T_*Z*_ = (1⁄*z*_1_, …, 1⁄*z*_*n*_). Our generalized definition of a polygenicity measure with LD corresponding to a function *f* takes the form

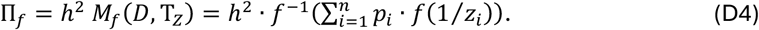

O’Connor et al.^21^ used an equivalent form for the harmonic polygenicity with LD.

This definition satisfies the intuitive requirements that we outlined at the beginning of this section. First, when casual variants are in LE it reduces to our previous definition of a polygenicity measure. Next, consider the case in which the LD matrix consists of blocks of perfectly linked variants with the blocks in LE with each other. Namely, the entries of the LD matrix 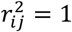 if (casual) variants *i* and *j* are in the same block and 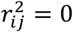 otherwise. We denote the blocks by indices *b* = 1, …, *m*, and the variants in block *b* by indices *j*_*b*_ = 1, …, *n*_*b*_, where 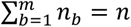. The expected marginal contribution of any variant in block *b* is therefore 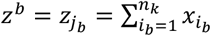 and the expected direct contribution of block *b* to heritability is 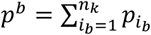. In this case, the polygenicity

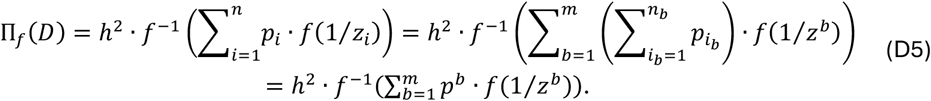

Namely, the polygenicity equals that of the effective variants representing blocks, where the effective variants are in LE. More generally, the way in which we weight causal variants assures that their causal effects are counted exactly once. Lastly, as we show in Appendix X, when variants are arranged in blocks with partial LD among them and LE among variants in different blocks, polygenicity measures take a value that is in between the case with block in perfect LD and the case with all variants in LE.

Consider the arithmetic polygenicity (*P* _*A*_) as an example. As we already saw, without LD, *P* _*A*_ counts the number of variants affecting the trait under consideration. In the general case, with any LD matrix, it takes the form

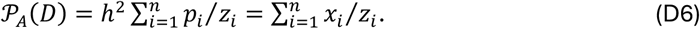

A causal variant in LE with all others has *z*_*i*_ = *x*_*i*_, and therefore contributes 1 to the count. Causal variants in a perfectly linked block that are in LE with all other causal variants jointly contribute 1 to the count. When two causal variants with direct contributions *x*_1_ = *x*_2_ = *x* are in partial LD with each other, with 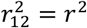, and are in LE with all other causal variants, their marginal contributions are *z*_1_ = *z*_2_ = (1 + *r*^2^)*x*. Their joint contribution to the count *x*_1_⁄*z*_1_ + *x*_2_⁄*z*_2_ = 2⁄(1 + *r*^2^) decreases monotonically with increasing *r*^2^, and is always between a contribution of 1 in the case of perfect LD and 2 in the case of LE.

## APPENDIX E estimating polygenicity using FMR

Polygenicity can be estimated using Fourier Mixture Regression (FMR)^24^, which fits a flexible approximation to the non-i.i.d. normal model that is defined in Appendix D. Specifically, FMR assumes that if variant *j* has causal effect-size variance *p*_*j*_ > 0, then its marginal effect-size variance, *z*_*j*_, is approximately one out of 13 discrete values, 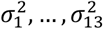. These values are chosen on the basis of the observed test statistics using a heuristic. FMR estimates the total fraction of heritability explained by variants of each effect size:

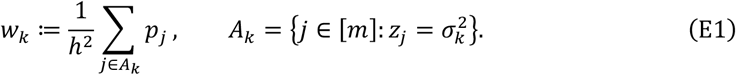

In principle, the resulting model fit can be used to estimate any polygenicity measure. Let 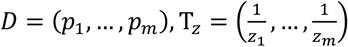, and 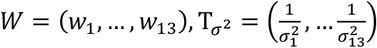. Then the estimand is related to the FMR model fit as follows:

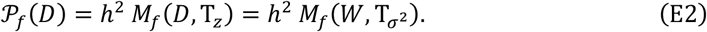

Substituting estimated for true values of *W* recovers the estimator (equation xy).

It can be necessary to implement the functions *f* and *f*^−1^ in a manner that is numerically stable; for example, the function

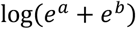

is implemented by calculating

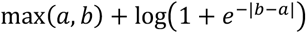

which avoids overflow when *e*^*a*^ or *e*^*b*^ is large. (This is used to implement the softmin function).

FMR estimation cannot work well for Π_*A*_ at finite sample size. If effect sizes can be arbitrarily small, then any fixed choice of mixture components will lack sufficiently small components. The formula in this special case becomes:

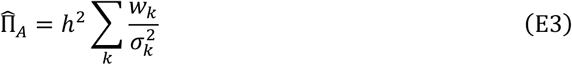

and if extremely small components 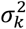 are included, then the ratio 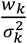 becomes unstable. In practice, estimates of Π_*A*_ continually become larger and noisier as smaller components are added.

More generally, FMR estimates of a polygenicity measure are expected to be consistent but not unbiased at finite sample size, and the behavior of the estimator can be highly dependent upon the measure that is chosen. We recommend using simulations to determine whether its estimates can be relied upon in practice. The same guidance applies to other estimation methods, as some measures of polygenicity (like *P*_*A*_) are fundamentally difficult to estimate via any method.

When analyzing real traits with FMR, we exclude the HLA region. This choice is necessary due to the HLA region’s unusual LD patterns, which extend beyond the map-distance boundaries used to define Fourier LD scores. If the region could be included, it would cause a reduction in our polygenicity estimates for autoimmune diseases (inflammatory bowel disease and hypothyroidism; a major cause of hypothyroidism is Hashimoto’s disease).

## APPENDIX F simulations

We performed simulations involving SNPs with MAF>0.01 in UK Biobank^42^ self-reported white individuals on chromosome 1. We simulated the data using LD graphical models and the *graphld* software, which allow computationally efficient simulation of summary statistics from their asymptotic sampling distribution with realistic patterns of local LD, as described in ref. ^43^. We performed 100 replicates. FMR was applied to the simulated summary statistics in conjunction with LD reference data from 1000 Genomes Europeans^44^ (which only approximately matches LD patterns in UKB). FMR was run with 13 Normal mixture components (its default settings^24^).

For the simulated effect size distribution, we randomly choose a fraction *q*_*L*_ of SNPs to be causal with ‘large’ direct contributions chosen from a Normal distribution 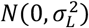, and a fraction *q*_*S*_ of SNPs to be causal with ‘small’ direct contributions chosen from a Normal distribution 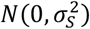, where 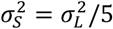. The remaining SNPs are taken to have no direct effect (i.e., to be non-causal). In our primary simulations (Figure 2), parameters were:

- 1/*q*_*S*_ = 200, 500, 1000, 2000, 5000, 10000
- *q*_*L*_ = *q*_*s*_/5
- *h*^2^ = 0.1
- 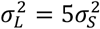, chosen such that 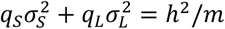 where *m* is the number of SNPs
- *α* = −1, i.e., no dependency of per-SNP heritability on allele frequency
- GWAS sample size *N* = 100,000

We performed three secondary simulations (Figure F1). First, we simulated a three-component model, using three mixture components with effect size variance 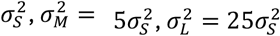. They had mixture weights:

- 1/*q*_*S*_ = 200, 500, 1000, 2000, 5000, 10000
- *q*_*M*_ = *q*_*s*_/5
- *q*_*L*_ = *q*_*s*_/25

and the value of *σ*^2^ was chosen such that 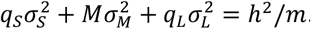.

Second, we performed a simulation at smaller sample size, reducing *N* from 100,000 to 20,000.

Third, we simulated allele frequency-dependent architecture, with an *α* parameter of −0.5. This causes low-frequency variants to have smaller contributions to heritability (but larger per-allele effect sizes). It was implemented by scaling all of the normalized effect size variance parameters by the factor (*pq*)^1+*α*^ and then re-normalizing them such that the total heritability was 0.1.

**Figure F1.**
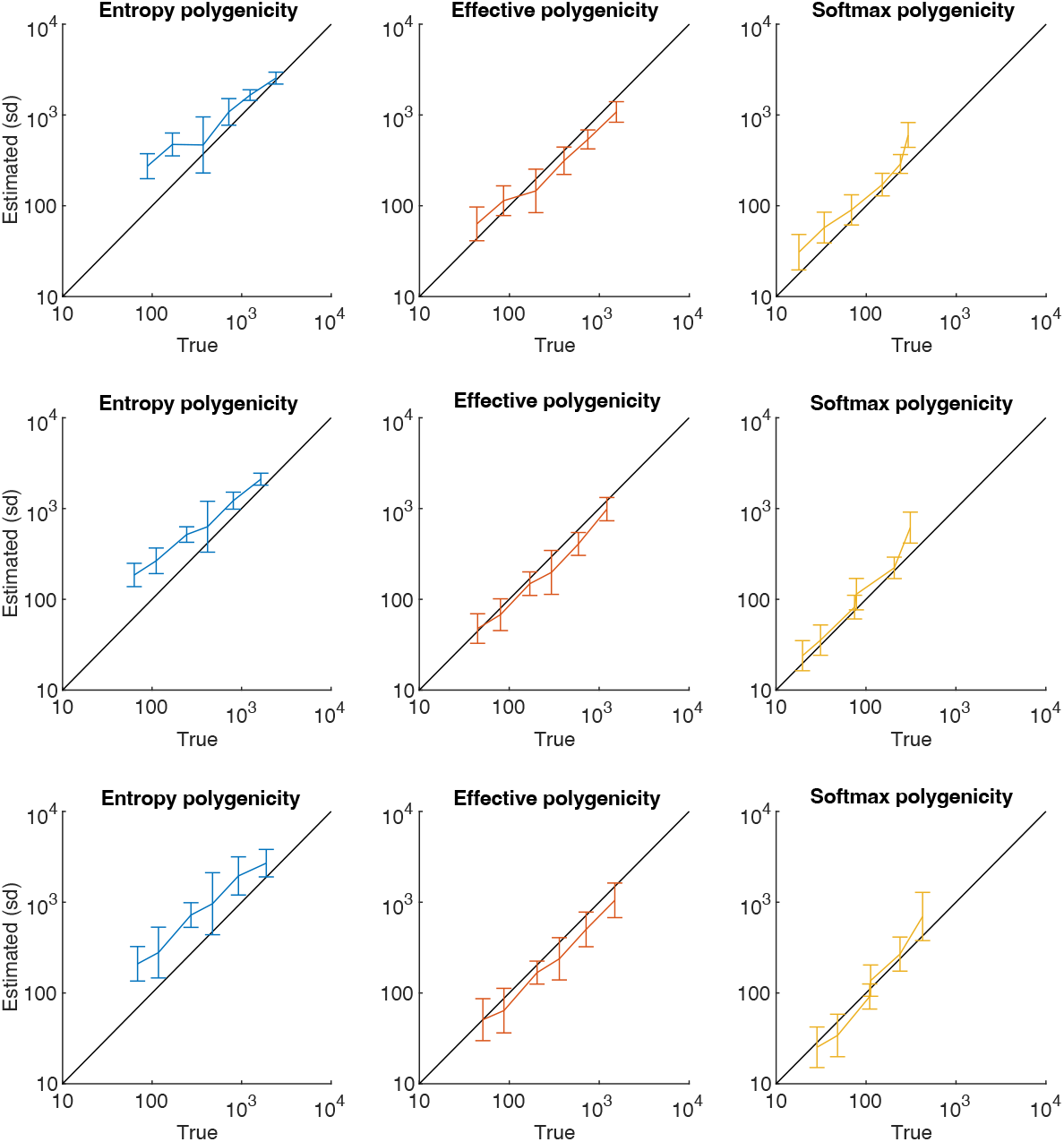
Entropy polygenicity of a Mendelian trait. See text for our assumptions. The graph was plotted based on Eq. G7, assuming *Lθ* = 1.

## APPENDIX G Entropy polygenicity of a Mendelian trait

Here we derive a rough approximation for the entropy polygenicity of a Mendelian trait. This is very much a back of the envelope calculation intended to clarify why we say that the polygenicity of Mendelian traits is much lower than the polygenicity of the traits we consider in Figure 3.

We assume a panmictic, diploid population of constant size *N* and that mutations affecting the trait arise at a single gene with *L* sites, with mutation rate *u* per site per gamete per generation. We further assume that *θ* = 4*Nu* ≪ 1, allowing us to apply the infinite sites approximation, and that variants are at LE with one another, allowing us to treat sites as independent. Lastly, we assume that all variants in the gene have the same effect size *a* and are subject to additive purifying selection with the same population scaled selection coefficient *γ*. This model is similar to the one in ref. ^45^.

We use the diffusion approximation to calculate the distribution of variant contributions to variance^46^. A variant with derived allele frequency (DAF) *x* contributes *v*(*x*) = 2*a*^2^*x*(1 − *x*) to variance. To calculate the expectations per site, we multiply the rate at which mutations arise by the expected total contribution of an individual mutation during its sojourn in the population. Namely,

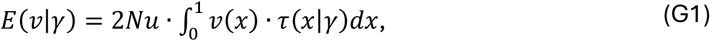

with the sojourn time

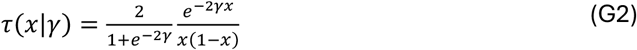

defined such that τ(*x*|*γ*)*dx* is the expected number of generations that an allele with scaled selection coefficient *γ* that originated from a mutation (with frequency 1⁄2*N*) spends between frequencies *x* and *x* + *dx*. This way we find that the expected contribution per site is

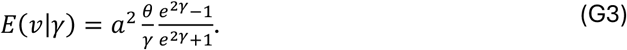

Next, we approximate the expected entropy polygenicity. The logarithm of this expectation can be approximated by

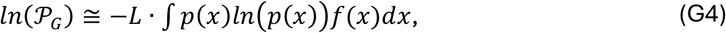

where

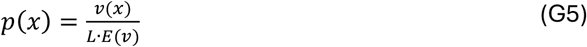

is the proportional contribution of a variant with DAF *x* to heritable variance, where we dropped the dependence on *γ* for simplicity (as it is held constant), and

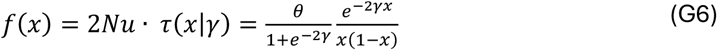

is the probability density of having a variant with DAF *x* at a site. After some algebra, we find that

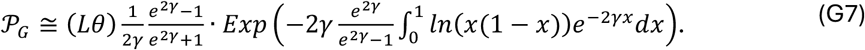

Figure G1 shows how the entropy polygenicity depends on the scaled selection coefficient. Here we have taken *Lθ* = 1; assuming that *θ* = 10^−3^—a typical value for human populations—this implies that *L* = 10^3^, which is a sensible, perhaps even high value for the number of sites in a gene that would affect a trait. Taking a greater value of *Lθ* would amount to multiplying the polygenicity by a corresponding fold increase. The graph illustrates that the maximal polygenicity is attained assuming that variants are selectively neutral (*γ* = 0). Even in that case, we find the entropy polygenicity to be <3.75, which is more than two orders of magnitude lower than the two lowest values we estimated and more than three orders of magnitude lower than the other 34/36 traits we examined.

**Figure G1.**
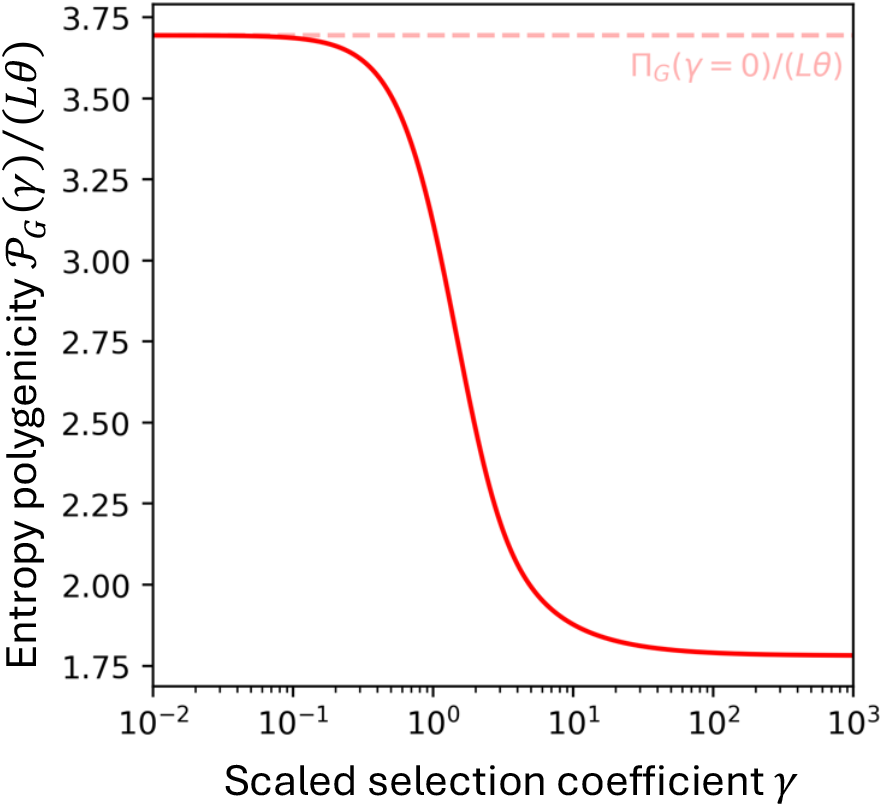
polygenicity estimates in simulations. (Top row) simulations involving an alternative effect-size distribution, with a mixture of three non-null Gaussian distributions. Each Gaussian distribution had an equal contribution to heritability, and their variances differed from each other by a factor of 5 (for a total difference of 25 between the small-and large-effect components). (Second row) simulations at smaller sample size (N=20k, vs. 100k in other simulations). (Third row) simulations with allele frequency dependent architecture, with an alpha parameter of ×0.5 (compared with ×1 for other simulations). Markers and error bars indicate means and standard deviations across 10 replicates. The same random seeds were used across the three simulations, so their sampling variability is correlated.

## Supplementary Table Captions

**Supplementary Table 1: Summary statistics included in the study**. We analyzed the 32 traits that were analyzed in ref. 24, together with four additional traits, all from UK Biobank (triglycerides, LDL cholesterol, HbA1c, and hair color). These traits were added because they seemed likely to have lower polygenicity.

**Supplementary Table 2: Polygenicity estimates for 36 traits**. For each phenotype, we report the point estimate and standard error of the log10-polygenicity for the three polygenicity measures. We also report mutational target size estimates from ref. 25 for traits that overlapped with those in our study, as well as the number of genome-wide significant lead SNPs as defined in ref. 24.

## References

1. Jobling, M., Hollox, E., Hurles, M., Kivisild T. C Tyler-Smith C. HUMAN EVOLUTIONARY GENETICS Second Edition. (2014).

2. Lango Allen, H. et al. Hundreds of variants clustered in genomic loci and biological pathways affect human height. Nature 467, 832–838 (2010).

3. International Schizophrenia Consortium et al. Common polygenic variation contributes to risk of schizophrenia and bipolar disorder. Nature 460, 748–752 (2009).

4. Falconer, D. S. C Mackay, T.F.C. Quantitative Genetics. (Longman London, 1983).

5. Visscher, P. M., Yengo, L., Cox, N. J. C Wray, N. R. Discovery and implications of polygenicity of common diseases. Science (1S7S) 373, 1468–1473 (2021).

6. Risch, N. et al. A genomic screen of autism: evidence for a multilocus etiology. The American Journal of Human Genetics 65, 493–507 (1999).

7. Yang, J. et al. Common SNPs explain a large proportion of the heritability for human height. Nat Genet 42, 565–569 (2010).

8. Wood, A. R. et al. Defining the role of common variation in the genomic and biological architecture of adult human height. Nat Genet 46, 1173–1186 (2014).

9. Bulik-Sullivan, B. K. et al. LD Score regression distinguishes confounding from polygenicity in genome-wide association studies. Nat Genet 47, 291 (2015).

10. Khera, A. V et al. Genome-wide polygenic scores for common diseases identify individuals with risk equivalent to monogenic mutations. Nat Genet 50, 1219 (2018).

11. Yengo, L. et al. A saturated map of common genetic variants associated with human height. Nature 610, 704–712 (2022).

12. Sinnott-Armstrong, N. et al. Genetics of 35 blood and urine biomarkers in the UK Biobank. Nat Genet 53, 185–194 (2021).

13. Zeng, J. et al. Signatures of negative selection in the genetic architecture of human complex traits. Nat Genet 50, 746–753 (2018).

14. Zhang, Y., Qi, G., Park, J.-H. C Chatterjee, N. Estimation of complex effect-size distributions using summary-level statistics from genome-wide association studies across 32 complex traits. Nat Genet 50, 1318 (2018).

15. Weissbrod, O. et al. Functionally informed fine-mapping and polygenic localization of complex trait heritability. Nat Genet 52, (2020).

16. Sinnott-Armstrong, N., Naqvi, S., Rivas, M. C Pritchard, J.K. GWAS of three molecular traits highlights core genes and pathways alongside a highly polygenic background. Elife 10, e58615 (2021).

17. Holland, D. et al. Beyond SNP heritability: Polygenicity and discoverability of phenotypes estimated with a univariate Gaussian mixture model. PLoS Genet 16, e1008612 (2020).

18. Stahl, E. A. et al. Bayesian inference analyses of the polygenic architecture of rheumatoid arthritis. Nat Genet 44, 483–489 (2012).

19. Palla, L. C Dudbridge, F. A Fast Method that Uses Polygenic Scores to Estimate the Variance Explained by Genome-wide Marker Panels and the Proportion of Variants Affecting a Trait. Am J Hum Genet G7, 250–259 (2015).

20. Moser, G. et al. Simultaneous discovery, estimation and prediction analysis of complex traits using a bayesian mixture model. PLoS Genet 11, e1004969 (2015).

21. O’Connor, L. J. et al. Extreme Polygenicity of Complex Traits Is Explained by Negative Selection. The American Journal of Human Genetics (2019).

22. Cover, T. M. Elements of Information Theory. (John Wiley C Sons, 1999).

23. Andrey, K. On the Notion of Mean. Mathematics and Mechanics 1GG, 144–146 (1930).

24. O’Connor, L. J. The distribution of common-variant effect sizes. Nat Genet 53, 1243– 1249 (2021).

25. Simons, Y. B., Mostafavi, H., Smith, C. J., Pritchard, J. K. C Sella, G. Simple scaling laws control the genetic architectures of human complex traits. bioRxiv 2010–2022 (2022).

26. Weiner, D. J. et al. Polygenic architecture of rare coding variation across 394,783 exomes. Nature 614, 492–499 (2023).

27. Kaplanis, J. et al. Evidence for 28 genetic disorders discovered by combining healthcare and research data. Nature 586, 757–762 (2020).

28. Simons, Y. B., Bullaughey, K., Hudson, R. R. C Sella, G. A population genetic interpretation of GWAS findings for human quantitative traits. PLoS Biol 16, e2002985 (2018).

29. Sella, G. C Barton, N.H. Thinking about the evolution of complex traits in the era of genome-wide association studies. Annu Rev Genomics Hum Genet 20, 461–493 (2019).

30. Spence, J. P. et al. Specificity, length, and luck: How genes are prioritized by rare and common variant association studies. bioRxiv 2012–2024 (2024).

31. Berg, J. J., Li, X., Riall, K., Hayward, L. K. C Sella, G. Mutation-selection-drift balance models of complex diseases. bioRxiv 2025 (2025).

32. Koch, E. et al. Genetic association data are broadly consistent with stabilizing selection shaping human common diseases and traits. bioRxiv 2024–2026 (2024).

33. Boyle, E. A., Li, Y. I. C Pritchard, J. K. An Expanded View of Complex Traits: From Polygenic to Omnigenic. Cell 16G, 1177–1186 (2017).

34. Liu, X., Li, Y. I. C Pritchard, J. K. Trans effects on gene expression can drive omnigenic inheritance. Cell 177, 1022–1034 (2019).

35. Lenormand, T. Gene flow and the limits to natural selection. Trends Ecol Evol 17, 183–189 (2002).

36. Yeaman, S. C Whitlock, M. C. The genetic architecture of adaptation under migration–selection balance. Evolution (N Y) 65, 1897–1911 (2011).

37. Kirkpatrick, M. C Barton, N. Chromosome inversions, local adaptation and speciation. Genetics 173, 419–434 (2006).

38. Yeaman, S. C Otto, S. P. Establishment and maintenance of adaptive genetic divergence under migration, selection, and drift. Evolution (N Y) 65, 2123–2129 (2011).

39. Yeaman, S. Evolution of polygenic traits under global vs local adaptation. Genetics 220, iyab134 (2022).

40. Crawford, N. G. et al. Loci associated with skin pigmentation identified in African populations. Science (1S7S) 358, eaan8433 (2017).

41. Nambiar, K. K., Varma, P. K. C Saroch, V. An axiomatic definition of shannon’s entropy. Appl Math Lett 5, 45–46 (1992).

42. Bycroft, C. et al. The UK Biobank resource with deep phenotyping and genomic data. Nature 562, 203–209 (2018).

43. Salehi Nowbandegani, P. et al. Extremely sparse models of linkage disequilibrium in ancestrally diverse association studies. Nat Genet 1–9 (2023).

44. Byrska-Bishop, M. et al. High coverage whole genome sequencing of the expanded 1000 Genomes Project cohort including 602 trios. bioRxiv 2021.02.06.430068 (2021) doi:10.1101/2021.02.06.430068.

45. Pritchard, J. K. Are rare variants responsible for susceptibility to complex diseases? The American Journal of Human Genetics 6G, 124–137 (2001).

46. Ewens, W. J. Mathematical Population Genetics. (No Title) (1964).

